# Molnupiravir inhibits Bourbon virus infection and disease-associated pathology in mice

**DOI:** 10.1101/2025.04.21.649883

**Authors:** Gayan Bamunuarachchi, Fang R. Zhao, Traci L. Bricker, Kuljeet Seehra, Andrew Karp, Michael S. Diamond, Adrianus C. M. Boon

## Abstract

Bourbon virus (BRBV) is an emerging tick-borne virus that can cause severe and fatal disease in humans. BRBV is vectored via the *Amblyomma americanum* tick, which is widely distributed throughout the central, eastern, and southern United States. Serosurveillance studies in Missouri and North Carolina identified BRBV-neutralizing antibodies in approximately 0.6% of tested individuals. To date, no specific antiviral therapy exists. Molnupiravir, an antiviral drug with oral bioavailability, has shown broad-spectrum antiviral activity against RNA viruses, including SARS-CoV-2. Here, we investigated the antiviral activity of molnupiravir against BRBV infection in cell culture and a mouse model of BRBV disease. Molnupiravir suppressed BRBV production in cells. *In vivo*, pre-exposure administration of molnupiravir protected susceptible type I interferon receptor knockout (*Ifnar1*^-/-^) mice against lethal BRBV infection. The protection by molnupiravir was associated with lower virus burden in mouse tissues, improvement of T cell (CD4^+^, CD8^+^) and B cell (follicular) profiles in the spleen, improvement of severe thrombocytopenia, and reduced pathology in the spleen and liver of BRBV-infected animals. Finally, therapeutic administration of molnupiravir starting 24 or 48 hours after infection ameliorated weight loss, clinical signs of disease, and lethality associated with BRBV infection. Overall, our experiments suggest that molnupiravir as a potential antiviral therapy for evaluation in humans with BRBV infections.

## INTRODUCTION

The incidence of emerging and reemerging viral infectious diseases has increased over the last 10 years, as evidenced by the COVID-19 pandemic in 2020, the numerous Ebola virus outbreaks in 2014-2020, the Zika virus epidemic in 2015, and the recent monkeypox outbreak. These outbreaks impact public health and can cause a substantial socioeconomic burden. Vector-borne diseases account for nearly 30% of emerging infectious diseases reported in the last decade [1]. As a result of climate change and deforestation, the rate of vector-borne infections, including tick-borne, is rising. Several tick species are endemic to the United States and carry human viral pathogens including Colorado tick fever virus, Powassan virus, Heartland virus, and Bourbon virus (BRBV).

BRBV was first identified in 2014 in the blood of a farmer from Bourbon County, Kansas, who reported a recent bite from a lone star tick (*Amblyomma americanum*) [2]. BRBV belongs to the family *Orthomyxoviridae,* genus *Thogotovirus,* and contains a segmented, negative sense, single-strand RNA genome. The six genome segments encode for membrane glycoprotein (GP), matrix protein (M), nucleoprotein (NP), and three viral polymerases (PB2, PB1, and PA). PB2, PB1, and PA form the RNA-dependent RNA polymerase (RdRp) complex [3] that, together with NP can generate messenger RNA and copy the genome. As BRBV is isolated predominantly from lone star ticks (*Amblyomma americanum)* [4–8], these arachnids are the likely competent vector of the virus. *Amblyomma americanum* is widely distributed across the Eastern, Southeastern, and Midwestern United States [9], suggesting that a large geographical area may be at risk of BRBV exposure.

Since its discovery in 2014, a total of five human BRBV cases have been reported, including a fatal infection in St. Louis County, Missouri in 2017 [10]. Although few human cases are documented, this may be due in part to low disease awareness among physicians and the lack of available diagnostic testing modalities. Indeed, two human serosurveillance studies, conducted in Missouri and North Carolina, identified BRBV-specific neutralizing antibodies in approximately 1 of 300 individuals [11, 12]. This suggests humans may be more at risk for exposure than suspected, including potential underdiagnosis of severe cases, and a need for antiviral countermeasures against BRBV.

Currently, there are no FDA-approved antiviral therapies for BRBV disease. Favipiravir, a viral RNA polymerase inhibitor that was developed as an antiviral drug targeting influenza A virus [13], showed efficacy against BRBV in a mouse model of infection [10]. Despite its pre-clinical efficacy against BRBV, favipiravir is not licensed in the USA, limiting its possible use. Several studies have evaluated antiviral therapeutics against BRBV in cell culture. Treatment with ribavirin, a known inhibitor of virus RdRp, showed antiviral activity against BRBV *in vitro* [14]. Myricetin, a parvovirus-nicking inhibitor, efficiently inhibited the RdRp activity of BRBV and virus replication in cells [15]. Recent high-throughput antiviral drug screening revealed that some dihydroorotate dehydrogenase inhibitors potently inhibited BRBV RdRp activity [16]. However, the lack of pre-clinical testing of these small molecule compounds has limited the development path for BRBV therapeutics. Repurposing previously approved antiviral drugs could overcome this barrier to the treatment of future BRBV cases. In this study, we evaluated several broad-spectrum antiviral compounds against BRBV in a pre-clinical setting and showed that molnupiravir, a nucleoside analog that has FDA Emergency Use Authorization (EUA) for SARS-CoV-2, effectively inhibits BRBV infection *in vitro* and *in vivo*.

## METHODS

### Ethics statement

All mouse experiments approved by the Institutional Animal Care and Use Committee (IACUC) at Washington University in St. Louis (Assurance # A3381-01).

### Virus and cell culture

A549 cells (gift from R. Webby, St. Jude Children’s Research Hospital) were maintained in DMEM media (Corning Cellgro) supplemented with 10% FBS (Biowest), 2 mM L-glutamine (Corning), 100 U/mL penicillin (Life Technologies), 100 μg/mL streptomycin (Life Technologies), and 1x MEM vitamins (Corning). Vero E6 cells (CRL-1586, ATCC) were grown in the same conditions above. HEK293 cells (gift from R. Webby, St. Jude Children’s Research Hospital) were grown in Opti-MEM (Corning Cellgro) media supplemented with 10% FBS (Biowest), 2 mM L-glutamine (Corning), 100 U/mL penicillin (Life Technologies), 100 μg/mL streptomycin (Life Technologies), and 1x MEM vitamins (Corning). All cells were maintained in a 37°C cell incubator with 5% CO_2_. Reverse genetics derived BRBV (strain BRBV-STL) [10], Dhori virus (DHOV, strain A/India/1313/1961), and thogotovirus (THOV, strain SiAr/126/72) were expanded on Vero E6 cells, aliquoted, stored at -80°C, and sequence verified prior to use in this study.

### In vitro antiviral activity testing

A549 cells (5 x 10^5^ cells/well) were seeded in 12-well plates and cultured overnight. Next, the cells were treated with different concentrations of remdesivir (Selleckchem; S8932), sofosbuvir (Selleckchem; S2794), N^4^-hydroxycytidine (NHC/EIDD-1931; Selleckchem; S0833), favipiravir (Selleckchem; S7975), or galidesivir (MedChemExpress; HY-18649/CS-3778) for 2 h in serum-free DMEM. Cells were then washed with PBS (1x) and inoculated with BRBV or DHOV at a multiplicity of infection (MOI) of 0.01 in a serum-free medium for 1 h. Next, the inoculation media was replaced with DMEM media containing 0.1% bovine serum albumin (BSA, Sigma) plus the different concentrations of antiviral compounds. At 24-, 48-, and 72-h post-inoculation (hpi), supernatants were collected, and the amount of infectious virus was quantified by focus focus-forming assay (FFA) for BRBV. Dimethyl sulfoxide (DMSO) was used as the control in the assay. In separate experiments, A549 cells were treated with different concentrations of NHC and subsequently inoculated with THOV or DHOV at a MOI of 0.01 in a serum-free medium, and virus yield at 48 hpi was measured by FFA or plaque assay, respectively.

### Virus titrations

FFA was used to quantify the amount of BRBV and THOV in the culture supernatant or tissues of infected mice. Monolayers of Vero E6 cells, seeded in 96 well plates, were washed once with PBS and incubated with three-fold dilutions of the cell supernatants or supernatants of homogenized tissues starting at a 1:10 dilution. After 1 h incubation at 37°C, the inoculum was aspirated, the cells were washed once with PBS, and 100 µL containing 1% methylcellulose (Sigma) and 1x minimum essential media (MEM, Corning) supplemented with 2% FBS was added to each well. The cells were incubated for 48 h at 37°C supplemented with 5% CO_2_. The monolayer was fixed with 100 μL of 5% formalin (Fisher Chemicals) for 30 min at room temperature. Next, cells were washed with PBS and subsequently stained with antibodies against the surface glycoprotein of BRBV or THOV. Subsequently, stained infected cells were further developed by using horseradish peroxidase (HRP) tagged secondary antibody as previously described [11]. Plaque assays were used to determine the amount of infectious DHOV in supernatants of cell culture or tissue homogenates. Briefly, Vero E6 (5 x 10^4^) cells were seeded into 24-well plates and cultured for 24 h. The cells were washed once with PBS and inoculated with serial ten-fold dilutions of the supernatant starting at 1:10 dilution. After 1 h at 37°C, the inoculum was aspirated, the cells were washed with PBS and overlaid with 1% methylcellulose and 1x MEM supplemented with 2% FBS. After 5 days, the cell monolayer was fixed with 5% formalin for 30 min at room temperature. Next, cells were washed with PBS and subsequently stained with 0.5% crystal violet for 1 h. The plaques were counted and used to determine the infectious titer as previously described [10].

### Viral RNA analysis

To quantify viral load in serum, lung, liver, and spleen tissue homogenates, RNA was extracted from 100 μL samples using E.Z.N.A. Total RNA Kit I (Omega) and eluted with 50 μL of water. Four microliters of eluate were used for real-time RT-qPCR to detect and quantify viral RNA (segment NP) of BRBV using TaqMan RNA-to-CT 1-Step Kit (Thermo Fisher Scientific) as previously described [17] using the following primers and probes: forward: GCAAGAAGAGGCCAGATTTC; reverse: TCGAATTCGGCATTCAGAGC; probe: CCTCACACCACGGAAGCTGGG; 5′Dye/3′Quencher: 6-FAM/ZEN/IBFQ. Viral RNA copy numbers per milliliter for tissue homogenates or serum are determined using a standard included in the assay. This standard was generated through *in vitro* transcription of a synthetic DNA molecule that contains the target region of the BRBV NP gene.

### Cell viability assay

The cytotoxicity of molnupiravir on A549 was evaluated by Cell Titer Blue® assay (Promega). Cells were seeded at 4 x 10^4^ cells per 100 µL in each well of a 96-well plate overnight. The next day, cells were washed and different concentrations of molnupiravir, starting at 500 µM and diluted in serum-free DMEM, were added to each well (final volume of 100 µL). After 48 h, cells were treated for 1 h at 37°C with Cell Titer Blue® reagent. The reduction of Resazurin to Resorufin in live cells was measured at 590 nm using a BioTek® plate reader and used to calculate the cell viability. Cells treated with 1% Triton X-100 were used as the positive control for cell death.

### BRBV polymerase activity assay

The BRBV polymerase activity assay was previously described [10]. Briefly, the BRBV firefly luciferase vector was generated by cloning the luciferase gene into the pLuci vector flanked by the 3’ and 5’ untranslated region of segment 6 of the original BRBV isolate (BRBV-KS). A total of 100 ng/per well of plasmid DNA split evenly between the 4 BRBV genes (PB1, PB2, PA, and NP) and the firefly reporter construct (firefly luciferase flanked by UTR’s of PB1), plus 5 ng of the Renilla luciferase transfection control was transfected into 293T cells in 96-well plates using TransIT LT1 (Mirus Bio). To test the effects of molnupiravir or favipiravir, various concentrations of the compound dissolved in DMSO were added to the well immediately after transfection. Firefly and renilla luciferase activities were measured 48 h after transfection using the Dual luciferase assay kit (Promega). The polymerase activity for each condition was reported as the average firefly/renilla (F/R) ratio of two wells per condition. Each assay and condition was repeated at least three independent times.

### Mouse experiments

Eight to ten-week-old male and female C57BL6/J or *Ifnar1*^-/-^ mice were used. *Ifnar1*^-/-^ mice were bred under specific pathogen-free conditions at Washington University and C57BL/6J mice were purchased from Jackson Laboratories (000664) and used for BRBV and DHOV studies, respectively. Toots and colleagues developed an isobutyric ester prodrug of NHC, molnupiravir, that increased its oral bioavailability in non-human primates [18]. Thus, we used molnupiravir in the mouse model. *Ifnar1*^-/-^ mice received molnupiravir (EIDD-2801, MedChemExpress; HY-135853/CS-0114880) at 50 mg/kg, 150 mg/kg, or 500 mg/kg in water containing 10% polyethylene glycol (Sigma) and 2.5% Cremophor® RH40 (Sigma) or vehicle only twice daily via oral gavage starting 4 h prior to or 24 and 48 h post-challenge with a lethal dose of BRBV. The drug treatment continued for 8 days or until the mice fully recovered their initial body weight, after the start of the treatment respectively. Weight change, clinical score (0 = normal, 1 = slightly ruffled fur, 2 = ruffled fur and hunched back, 3 = ruffled fur, hunched back, reduced activity, 4 = ruffled fur, hunched back, reduced activity and labored breathing, 5 = death or lost more than 25% of their body weight), and survival were monitored for 14 days after BRBV infection. Lung, spleen, blood, serum, and liver were collected at 3- or 6-days post-infection (dpi) to measure virus titer by RT-qPCR and infectious virus assay, for histology, and flow cytometry. C57BL/6J mice were used to assess the efficacy of molnupiravir against DHOV. Mice received a 500 mg/kg dose of molnupiravir 4 h prior to challenge with 10X mouse lethal dose 50 [MLD_50_ ∼ 1 plaque forming unit (pfu MLD_50_) of DHOV. Subsequently, drug treatment continued for 8 days (twice daily), and weight change, clinical signs were monitored.

### Histological analyses

Liver and spleen were collected from the vehicle control or molnupiravir-treated mice at 3 and 6 dpi, fixed in 4% PFA at room temperature for a week, and then dehydrated in 70% ethanol. Paraffin-embedded tissues were sectioned and stained with hematoxylin and eosin following standard procedures. Histological analyses were performed in a blinded manner. The severity score of liver pathology was determined based on the following criteria: inflammation score 0 = absent, score 1 = scattered, score 2 = foci, score 3 = diffuse; necrosis score 0 = absent, score 1 = < 10%, score 2 = 10-50%, score 3 = > 50%; and steatosis score 0 = < 10%, score 1 = 10-30%, score 2 = 31-60%, score 3 = > 60%. Spleen histopathology was scored based on the following: splenic disorganization score 0 = none, 1 = mild loss of white pulp distinction with blurring of marginal zone, 2 = loss of marginal zone, 3 = complete loss of structure; lymphoid depletion score 0 = none, 1 = mild, 2 = moderate, 3 = severe; and necrosis/ apoptosis score 0 = none, 1 = mild, 2 = moderate, 3 = severe.

### Flow cytometric analyses

At 6 dpi, whole blood from vehicle- or molnupiravir-treated mice was collected via cardiac puncture into EDTA tubes. To determine complete blood counts, samples of whole blood were incubated with Fc Block (BD Pharmingen) and then stained with a fixable live/dead dye (eBioscience) and the following fluorophore-conjugated antibodies in PBS with 2% FBS: BUV395 anti-mouse CD45 (RRID: AB_2739420), APC anti-mouse CD41 (RRID: AB_11126751), and PE anti-mouse TER-119 (RRID: AB_313709). For evaluation of circulating immune cell populations in the blood, erythrocytes were lysed with ACK Lysing Buffer (Gibco) before staining of remaining cells with fluorophore-conjugated antibodies. For evaluation of splenocytes, not including dendritic cells and macrophages, single-cell suspensions were obtained by mechanical dissociation of spleens, followed by passage through 70 µm nylon mesh filters and lysis of erythrocytes. The following additional antibodies were used in this study: AF700 anti-mouse CD11b (RRID: AB_493705), BV711 anti-mouse Ly6C (RRID: AB_2562630), BV785 anti-mouse Ly6G (RRID: AB_2566317), PE anti-mouse Siglec-F (RRID: AB_2750235), AF488 anti-mouse TCRb (RRID: AB_493344), APC-Cy7 anti-mouse CD19 (RRID: AB_830707), PE-Cy7 anti-mouse NK1.1 (RRID: AB_389364), PE-Cy5 anti-mouse NK1.1 (RRID: AB_493590), PE-Dazzle 594 anti-mouse CD4 (RRID: AB_2563685), PE-Cy5 anti-mouse CD8 (RRID: AB_312749), BV605 anti-mouse CD138 (RRID: AB_2562336), BUV395 anti-mouse CD19 (RRID: AB_2722495), BV711 anti-mouse B220 (RRID: AB_2563491), PE-Cy7 anti-mouse CD21/CD35 (RRID: AB_1953277), BV421 anti-mouse CD23 (RRID: AB_2563599), BV605 anti-mouse IgD (RRID: AB_2562887), and FITC anti-mouse IgM (RRID: AB_315056). Flow cytometry cell counting beads were used to quantitate absolute cell counts in the blood and spleen. All samples were processed on a BD LSRFortessa X-20 flow cytometer (BD Biosciences) and analyzed using FlowJo software (version 10.10.0).

### Statistical analysis

Statistical analyses were performed using GraphPad Prism 10.1 software. A minimum of three independent experiments were used to analyze *in vitro* data, and more than six animals per group were used for *in vivo* experiments. Mann-Whitney test, Kruskal-Wallis test, student’s t-test, and one-way ANOVA, followed by post hoc comparisons (Bonferroni correction, Dunnett’s, Sidak’s, and Tukey’s pairwise comparisons), and area under the curve analysis were used to compare two groups or multiple groups. Specific tests were selected based on the variance of the data and the number of comparator groups. A *P-*value of< 0.05 was considered significant.

### Data availability statement

All data needed to evaluate the conclusions in the paper are present in the paper and/or the Supplementary Materials or available online (DOI pending). This paper does not include the original code.

## RESULTS

### Antiviral screening of selected nucleoside analogs

We hypothesized that nucleoside analogs that showed broad-spectrum antiviral activity against other RNA viruses might inhibit BRBV infection. As such, we tested multiple compounds for activity against BRBV in a multi-cycle virus growth assay, including galidesivir, a potent RNA-dependent RNA polymerase inhibitor of influenza A virus [19]; molnupiravir and remdesivir, used to treat SARS-CoV-2 [20, 21]; and sofosbuvir, a nucleoside analog with activity against hepatitis C virus. Favipiravir, a known BRBV inhibitor, was included as a positive control, and virus accumulation in the supernatant in the presence or absence of the compounds was measured at 12, 24, 48, and 72 hpi by FFA. The biologically active form of molnupiravir, *N*^4^-hydroxycytidin (NHC, EIDD-1931), significantly reduced BRBV replication at 10 µM (*P* < 0.01) and 100 µM (*P* < 0.001) concentrations (**Fig. 1A**). At these doses of molnupiravir, the viability of the A549 cells was minimally affected (**Fig. S1**). Galidesivir also reduced virus titers, but only at 100 µM (*P* < 0.01), and breakthrough replication was observed (**Fig. 1B**). Antiviral activity against BRBV was not observed for remdesivir and sofosbuvir at concentrations ranging from 0.01 – 100 µM (**Fig. 1C-D**). The effective concentration (EC_50_) of molnupiravir (∼20 µM, **Fig. 1E**) was comparable to that of favipiravir (∼70 µM, **Fig. 1E and S2**).

**Figure 1.**
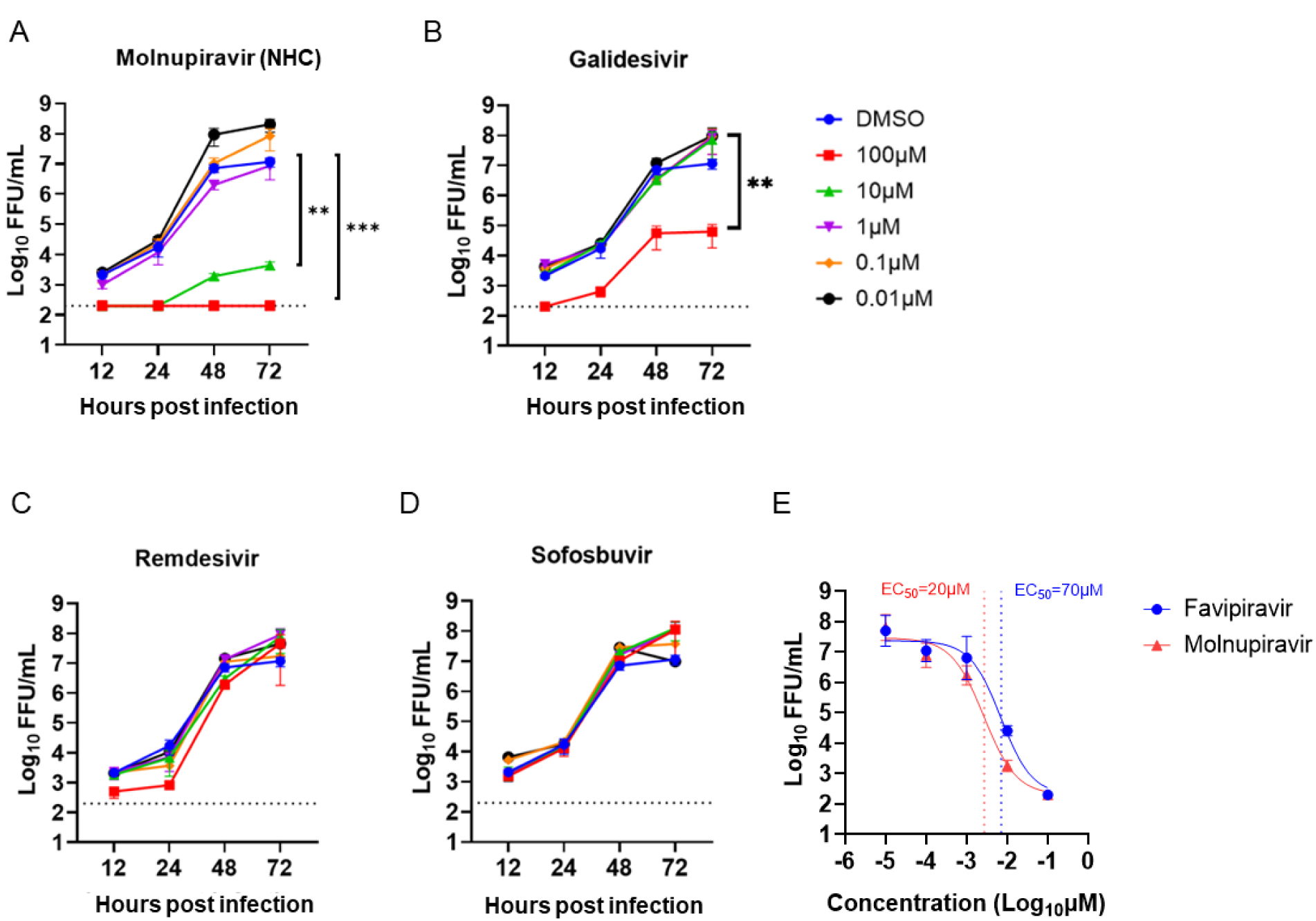
Molnupiravir inhibits BRBV infection in cell culture. (**A-D**) A549 cells inoculated with BRBV (MOI of 0.01) in the presence or absence of different concentrations of (**A**) molnupiravir (NHC), (**B**) galidesivir, (**C**) remdesivir, (**D**) sofosbuvir. The virus titer in the culture supernatant was quantified in 12, 24, 48, and 72 h. Values are means (± standard error of the mean) of the virus titer from three experiments performed in duplicate. ** *P <* 0.01, *** *P <* 0.005 by one-way ANOVA on the area under the curves fit by linear regression of log-transformed virus titer over time and compared with Dunnett’s multiple comparisons. The dotted line represents the limit of detection at 200 FFU/mL. (**E**) Inhibition comparison of molnupiravir and favipiravir in A549 cells at 48 hpi including indicated 50% effective concentration (EC_50_). A curve was fitted through the data using the log(inhibitor) vs. response-variable slope equation (GraphPad Prism 10.0).

### Prophylactic administration of molnupiravir protects mice from lethal BRBV infection

In contrast to wild-type C57BL/6J mice, congenic *Ifnar1*^-/-^ mice are highly susceptible to BRBV with 100% lethality after infection with 10-100 infectious units. BRBV infection results in liver damage, like that seen in BRBV-infected patients [10, 14]. To determine whether molnupiravir (EIDD-2801) can protect against lethal BRBV infection *in vivo*, we administered to male and female *Ifnar1*^-/-^ mice low (50 mg/kg), medium (150 mg/kg) or high (500 mg/kg) doses of molnupiravir or vehicle control via oral gavage 4 h prior to lethal challenge with BRBV (400X mouse lethal dose [MLD_50_ ∼ 1 plaque forming unit (pfu)]). Because the half-life of molnupiravir in mice is poor, we used higher dosing (50-500 mg/kg) schemes compared to other compounds; such doses are well tolerated in mice [22, 23]. Following BRBV infection, the mice continued to receive molnupiravir or the vehicle twice daily for 8 days. Body weight loss, clinical signs, and survival of the mice were monitored for 14 days. All three doses of molnupiravir significantly reduced weight loss and clinical disease scores in BRBV-infected mice compared to the vehicle control, with protective effects observed in a dose-dependent manner (**Fig. 2A-B**). The reduction in weight loss and clinical score of the high- and medium-dose treated *Ifnar1*^-/-^ mice was associated with uniform survival (*P* < 0.0001) after infection (**Fig. 2C and S3**). Low-dose molnupiravir-treated mice also showed improved survival (55%, *P* < 0.001) after infection (**Fig. 2C**).

**Figure 2.**
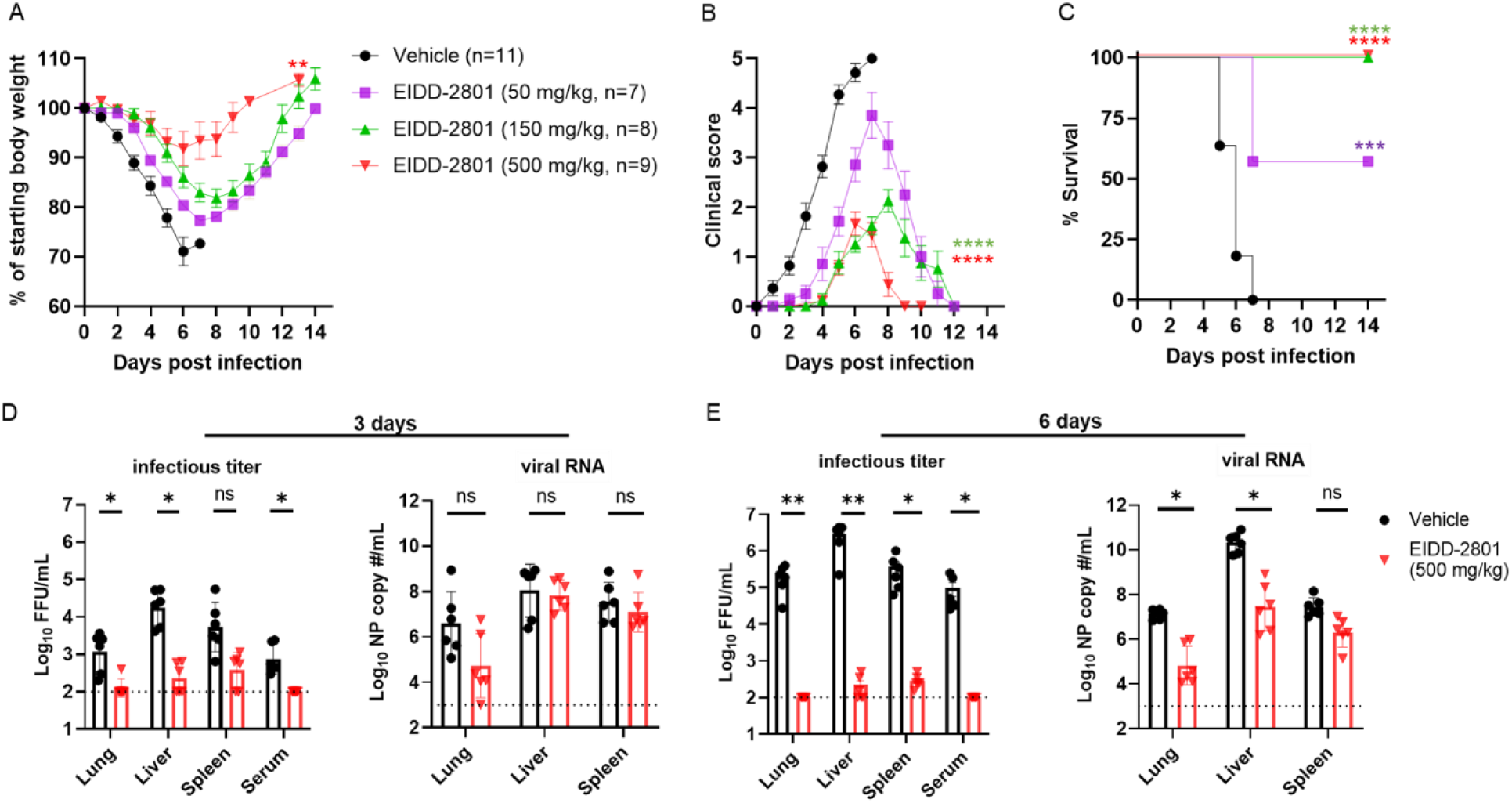
Prophylaxis with molnupiravir protects mice from fatal BRBV infection. (**A-C**) *Ifnar1*^-/-^ mice were treated with 50, 150, or 500 mg/kg of molnupiravir or vehicle control twice daily per oral gavage for eight days starting 4 h prior to inoculation with 400X MLD_50_ of BRBV. Weight change (**A**), clinical score (**B**), and survival rates (**C**) were monitored for 14 days. Values are the means (± standard error of the mean) of the groups. Weight change and clinical scores are analyzed by the one-way ANOVA followed by Dunnett’s multiple comparisons, and survival is analyzed by the log-rank test. ** *P <* 0.01; **** *P <* 0.0001. (**D-E**) *Ifnar1*^-/-^ mice were treated with 500 mg/kg of molnupiravir for 4 h prior to infection with 100X MLD_50_ of BRBV, and infectious virus titer and viral RNA levels at 3 or 6 dpi in different organs were measured by titration and RT-qPCR, respectively. Each data point represents a single mouse (n = 6 per group), and values are the means (± standard error of the mean) of the groups. Drug and vehicle-treated groups were analyzed by one-way ANOVA followed by Tukey’s post-test. * *P <* 0.05; ** *P* < 0.01.

Next, we measured the viral burden in the lung, liver, spleen, and serum of vehicle- and drug-treated (500 mg/kg) *Ifnar1*^-/-^ mice at 3 or 6 dpi with 100X MLD_50_ of BRBV. Virus titers were quantified by infectious virus assays (FFA) and RT-qPCR. Prophylaxis of molnupiravir reduced infectious BRBV levels in the lung (12-fold, *P* < 0.05), liver (85-fold, *P* < 0.05), spleen (25-fold, *P =* 0.09), and serum (11-fold, *P* < 0.05) at 3 dpi (**Fig. 2D**). Similarly, the amount of BRBV RNA was reduced in the lungs (126-fold) of the drug-treated mice, albeit the difference was not statistically significant. Differences in viral RNA levels were not detected in the liver or spleen of vehicle- and drug-treated mice at 3 dpi (**Fig. 2D**). At 6 dpi, the amount of infectious virus was reduced ∼2,100-fold (*P* < 0.01), ∼13,500-fold (*P* < 0.01), ∼1,250-fold (*P* < 0.05), and ∼950-fold (*P* < 0.05) in the lung, liver, spleen, and serum, respectively, of drug-treated mice (**Fig. 2E**). Additionally, the amount of BRBV RNA was significantly reduced in the lung (∼49-fold, *P* < 0.05) and liver (∼170-fold, *P* < 0.05), but not in the spleen (∼11-fold, *P* = 0.05) of drug-treated *Ifnar1*^-/-^ mice 6 dpi (**Fig. 2E**). These results show that molnupiravir can inhibit BRBV infection *in vivo* and reduce severe and fatal disease in immunodeficient mice.

### Therapeutic administration of molnupiravir protects *Ifnar1*^-/-^ mice from lethal BRBV infection

To determine if molnupiravir has therapeutic efficacy after infection, *Ifnar1^-/-^* mice were inoculated with 100X MLD_50_ of BRBV, and subsequently treated at either 1 or 2 dpi with a high dose (500 mg/kg) of molnupiravir twice daily until they fully recovered their initial body weight. Compared to vehicle control-treated mice, mice that received molnupiravir one day after a lethal dose of BRBV fully restored their body weight 14 days later, had a significantly (*P* < 0.01) lower clinical score, and survived (*P* < 0.01) infection (**Fig. 3A-C**). Initiating molnupiravir treatment 2 days after infection also reduced weight loss (*P* < 0.05) and clinical score (*P* < 0.05), and increased survival rates (*P* < 0.05), albeit with a smaller effect than animals receiving the drug 1 day after BRBV infection (**Fig. 3A-C**). Thus, molnupiravir treatment has both prophylactic and therapeutic efficacy against BRBV in a highly susceptible mouse model of infection and disease.

**Figure 3.**
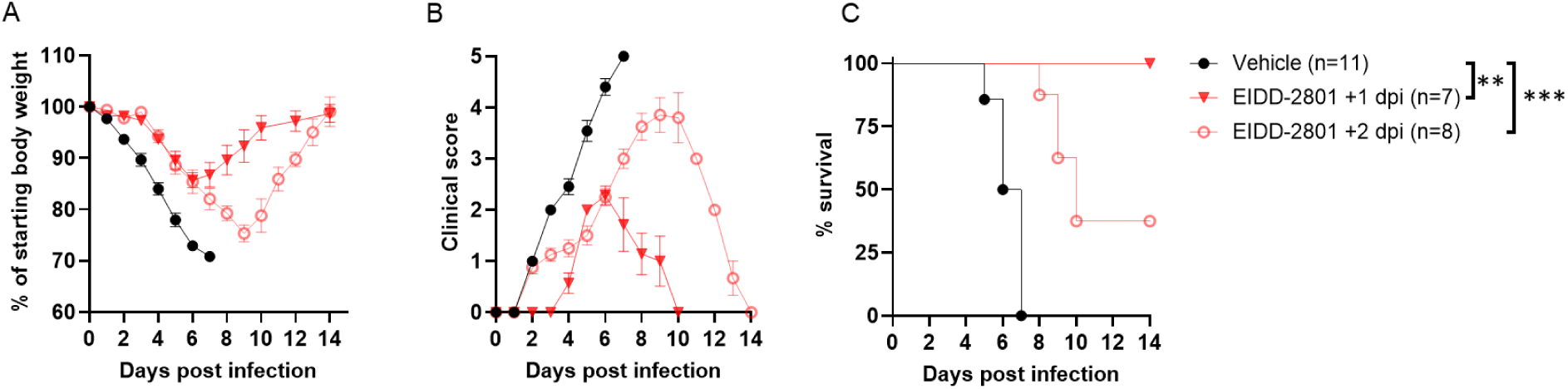
Therapeutic administration of molnupiravir protects mice from fatal BRBV infection. (**A-C**) *Ifnar1*^-/-^ mice were inoculated with 100X MLD_50_ of BRBV (IP) and treated with 500 mg/kg of molnupiravir (orally, twice daily) starting at one day (+1 dpi, red closed triangles, n = 7) or two days (+2 dpi, red open circles, n = 8) after infection until they fully recovered (reinstate the initial BW). Mice that received water containing 10% polyethylene glycol and 2.5% Cremophor® RH40 starting two days after inoculation (vehicle +2 dpi) were used as vehicle controls (n = 11). Weight change (**A**), clinical score (**B**), and survival rates (**C**) were monitored for 14 days. Values are the means (± standard error of the mean) and are analyzed by One-way ANOVA followed by Dunnett’s multiple comparison and survival by the log-rank test. ** *P <* 0.01; *** *P* < 0.005.

### Molnupiravir limits disease pathology in the spleen and liver of BRBV-infected mice

We next assessed the effect of molnupiravir on disease pathology in the spleen and liver of BRBV-infected mice. *Ifnar1^-/-^* mice developed splenomegaly after BRBV infection, with increased tissue weight at 6 dpi that improved with molnupiravir in prophylaxis treatment (*P* < 0.01, **Fig. 4A**). At a histological level, we observed disrupted splenic architecture in vehicle control-treated mice at 3 dpi, including effacement of the marginal zone and reduced numbers of lymphocytes in the periarteriolar lymphoid sheaths of the white pulp (**Fig. 4B-C**). Additionally, we detected foci of extramedullary hematopoiesis in the red pulp, as well as pigment-laden macrophages diffusely in both the red and white pulp (**Fig. 4B-C**). *Ifnar1^-/-^* mice treated high dose of molnupiravir (500 mg/kg) showed improvements in splenic architecture and lymphocyte numbers compared to vehicle-treated mice (*P* < 0.05, **Fig. 4B-C**). Although extramedullary hematopoiesis and pigment-laden macrophages in the spleen were still apparent at 3 dpi, by 6 dpi, vehicle control-treated mice showed almost complete loss of splenic architecture with diffuse mixing of the red and white pulps, substantially reduced numbers of lymphocytes, and necrosis (**Fig. 4A-B**). Although splenic pathology was still apparent, BRBV-infected *Ifnar1*^-/-^ mice treated with molnupiravir, showed marked improvement in overall architectural features (**Fig. 4B-C**).

**Figure 4.**
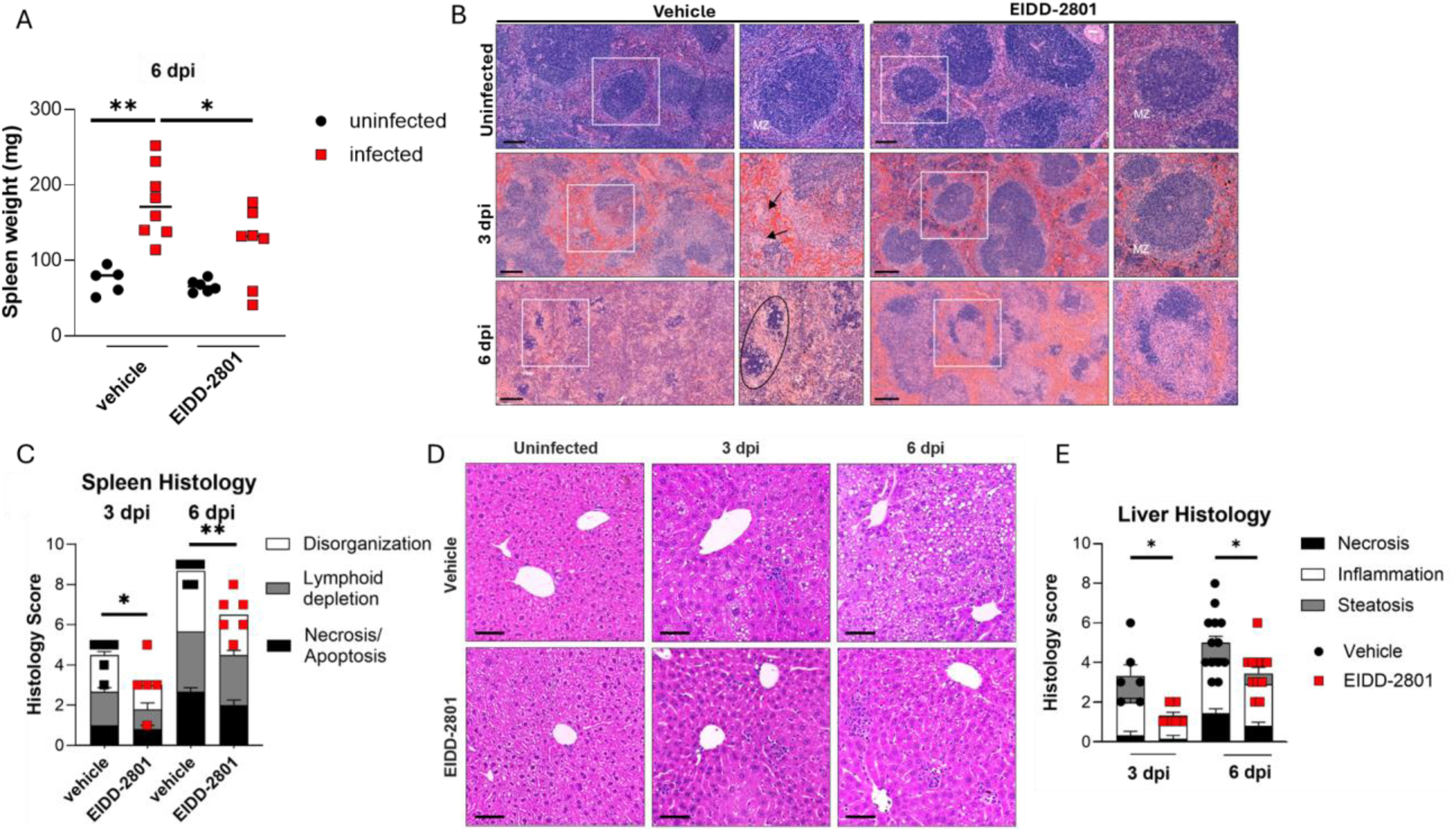
Molnupiravir reduces BRBV-associated pathology in the spleen and liver. *Ifnar1*^-/-^ mice were treated with either vehicle or 500 mg/kg of molnupiravir at 4 h prior to inoculation with 100X MLD_50_ of BRBV. Mice continued to receive 500 mg/kg of molnupiravir (orally, twice daily) until the day of harvest. (**A**) The gross weight of spleens from vehicle- or EIDD-2801-treated mice at 6 dpi. (**B-C**) Representative images of hematoxylin and eosin (H&E) staining of spleens from uninfected (n = 5 per group) or BRBV-infected (3 or 6 dpi, n = 5-6 per group) mice receiving vehicle or EIDD-2801 treatment (**B**) and scoring of spleen pathology (**C**). Circled regions show apoptotic bodies and tingible body macrophages with engulfed debris. MZ, marginal zone. Scale bar, 200 μm. (**D-E**) Representative images after H&E staining of livers from the vehicle or drug-treated mice that were uninfected (n = 5 per group) or BRBV-infected (3 or 6 dpi, n = 6-13 per group) (**D**) and scoring of liver pathology (**E**). Scale bar, 100 μm. In **A**, **C, and E**, each data point is from a single mouse, and bars represent the means (± standard error of the mean) of the groups. For analysis of spleen weight, spleen histology and liver histology of drug and vehicle-treated groups were from 2-3 independent experiments and analyzed by one-way ANOVA followed by Šidák’s post-test. * *P <* 0.05; ** *P <* 0.01.

Consistent with prior reports of liver injury in mice (*Ifnar1^-/-^*or *Stat1^-/-^*) and humans [14], we found that BRBV infection led to an accumulation of periportal and lobular inflammatory infiltrates, and steatosis (abnormal lipid retention), with minimal necrosis in vehicle control-treated mice at 3 dpi (**Fig. 4D-E**). By 6 dpi, BRBV-infected, control-treated mice had extensive inflammatory-induced necrosis of the liver cells (**Fig. 4D-E**). In molnupiravir-treated mice, liver pathology was improved at 3 and 6 dpi (*P* < 0.05) (**Fig. 4D-E**). matching the reduced viral burden in the liver at these time points (**Fig. 2D-E)**.

### Molnupiravir ameliorates splenic immune cell cytopenia and peripheral blood thrombocytopenia in BRBV-infected mice

BRBV infection in humans causes leukopenia, progressive thrombocytopenia, and lymphopenia [10]. Given the role of the spleen as a reticuloendothelial organ and that molnupiravir rescues BRBV-associated pathology (**Fig. 4**), we used flow cytometry to profile splenocyte populations from vehicle control- and molnupiravir-prophylactically treated mice at 6 dpi. Viability of splenocytes after BRBV infection, as detected by a fixable live/dead dye, was reduced in vehicle-treated mice and improved in mice treated with molnupiravir (**Fig. 5A**). Within the lymphoid population, BRBV infection led to reduced percentages and numbers of CD4^+^ (*P* < 0.001) and CD8^+^ (*P* < 0.01) T cells (**Fig. 5B-C**). This decline was partially rescued in mice treated with molnupiravir (**Fig. 5B-C**). B cells migrate from the bone marrow to secondary lymphoid organs, including the spleen, where they mature into follicular or marginal zone B cells. Consistent with our histological observations of splenic disorganization, vehicle-treated mice exhibited a marked reduction in both percentages (*P* < 0.001) and numbers (*P* < 0.001) of follicular and marginal zone B cells at 6 dpi; treatment with molnupiravir improved the numbers of follicular (*P* < 0.05) but not marginal zone (*P* = 0.10) B cells (**Fig. 5D-F**).

**Figure 5.**
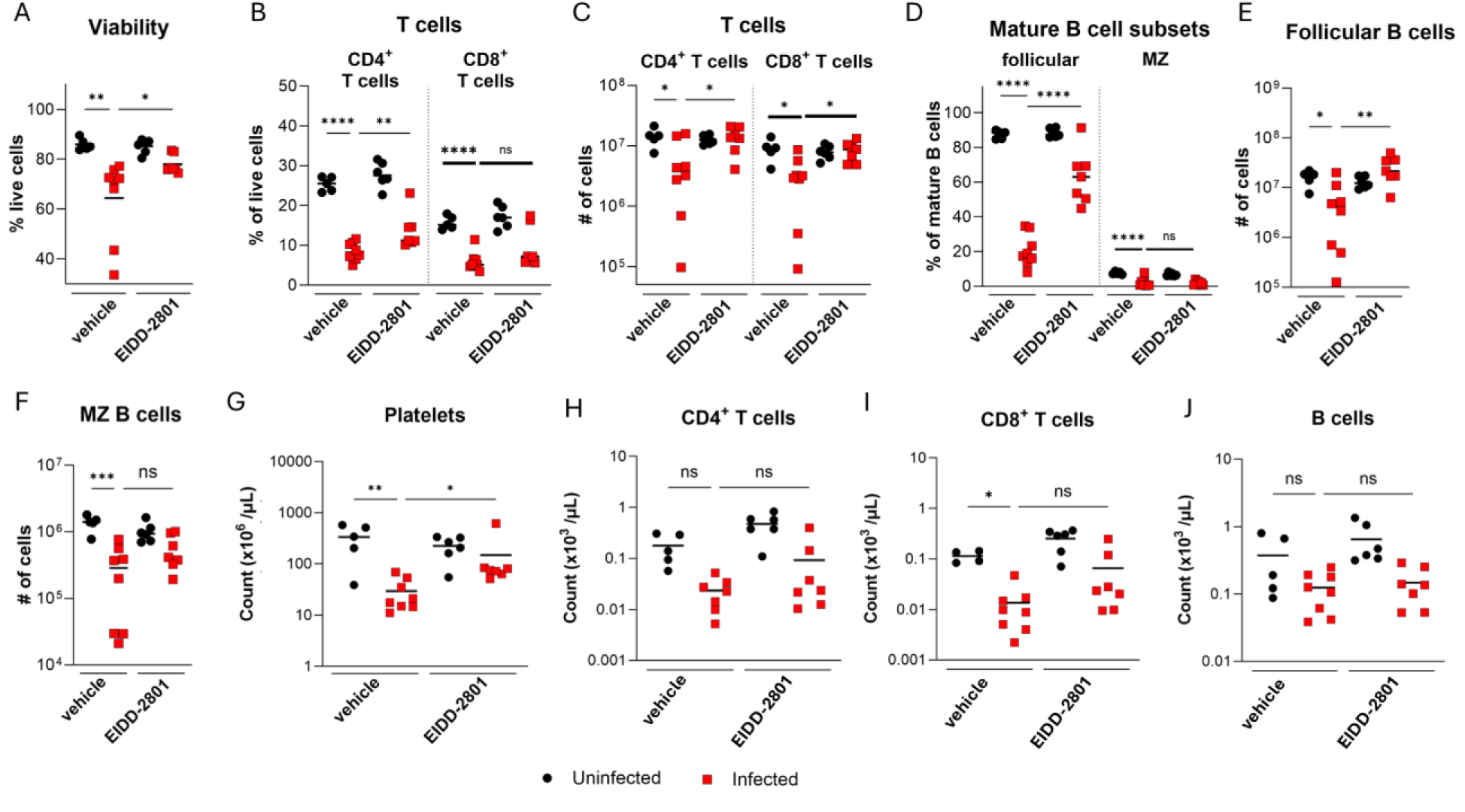
Molnupiravir improves BRBV-associated cytopenia in the spleen and peripheral blood thrombocytopenia. *Ifnar1*^-/-^ mice were treated with 500 mg/kg of molnupiravir at 4 h prior to inoculation with 100X MLD_50_ of BRBV, and the two-dose per day regiment was continued until day 6. Mouse splenocytes and blood were harvested at 6 dpi and evaluated by flow cytometry. (**A**) Viability of total splenocytes from vehicle- or EIDD-2801-treated mice at 6 dpi, as assessed by staining with a viability dye. (**B-C**) Percentages (**B**) and numbers (**C**) of splenic CD4^+^ and CD8^+^ T cells. (**D-F**) Percentages (**D**), and numbers of follicular (**E**) and marginal zone (MZ, **F**) B cells in the spleen. (**G-J**) Total counts of peripheral blood platelets (**G**), CD4^+^ T cells (**H**), CD8^+^ T cells (**I**), and B cells (**J**). Each data point represents a single mouse, and the bars represent the means of the groups. Drug and vehicle-treated groups were analyzed by one-way ANOVA with Šidák’s post-test. * *P <* 0.05; ** *P <* 0.01; *** *P <* 0.005; *** *P <* 0.005. The data are pooled from two independent experiments (n = 5-8 per group).

As BRBV infection in humans is known to cause progressive and severe thrombocytopenia [10], we investigated whether BRBV infection led to a similar cytopenia in mice, and molnupiravir treatment could reverse this phenotype. Like BRBV-infected humans, *Ifnar1^-/-^* mice became severely thrombocytopenic by 6 dpi (**Fig. 5G**), with mild declines in immune cell counts due primarily to T cell lymphopenia (**Fig. 5H-J**). Treatment of mice with molnupiravir improved platelet counts (*P* < 0.05, **Fig. 5G**), but not peripheral T or B cell numbers in the blood (**Fig. 5H-J**). Increased platelet counts in molnupiravir-treated mice suggest that the improvement might be due to reduced splenomegaly and/or improved liver function **(Fig. 4A, D-E)**.

### The antiviral mechanism of molnupiravir is not associated with direct inhibition of BRBV RdRp

While many nucleoside analogs directly inhibit the RNA-dependent RNA polymerase (RdRp) function of RNA viruses and viral genome replication [24], molnupiravir is thought to act as a mutagen and introduce fatal mutations in the genome of affected viruses [25]. To evaluate its potency against BRBV RdRp inhibition, we applied an established BRBV polymerase activity assay. At concentrations of molnupiravir that inhibit BRBV replication in A549 cells (**Fig. 1A**), polymerase activity was unaffected (**Fig. 6A**). In contrast, favipiravir did inhibit the polymerase activity of BRBV (**Fig. S4**). To confirm that molnupiravir was also effective in 293T cells, multi-cycle viral growth assays were repeated in these cells. Like A549 cells, significant reductions in BRBV titer at 10 and 100 µM concentrations of molnupiravir were detected (**Fig. 6B**).

**Figure 6.**
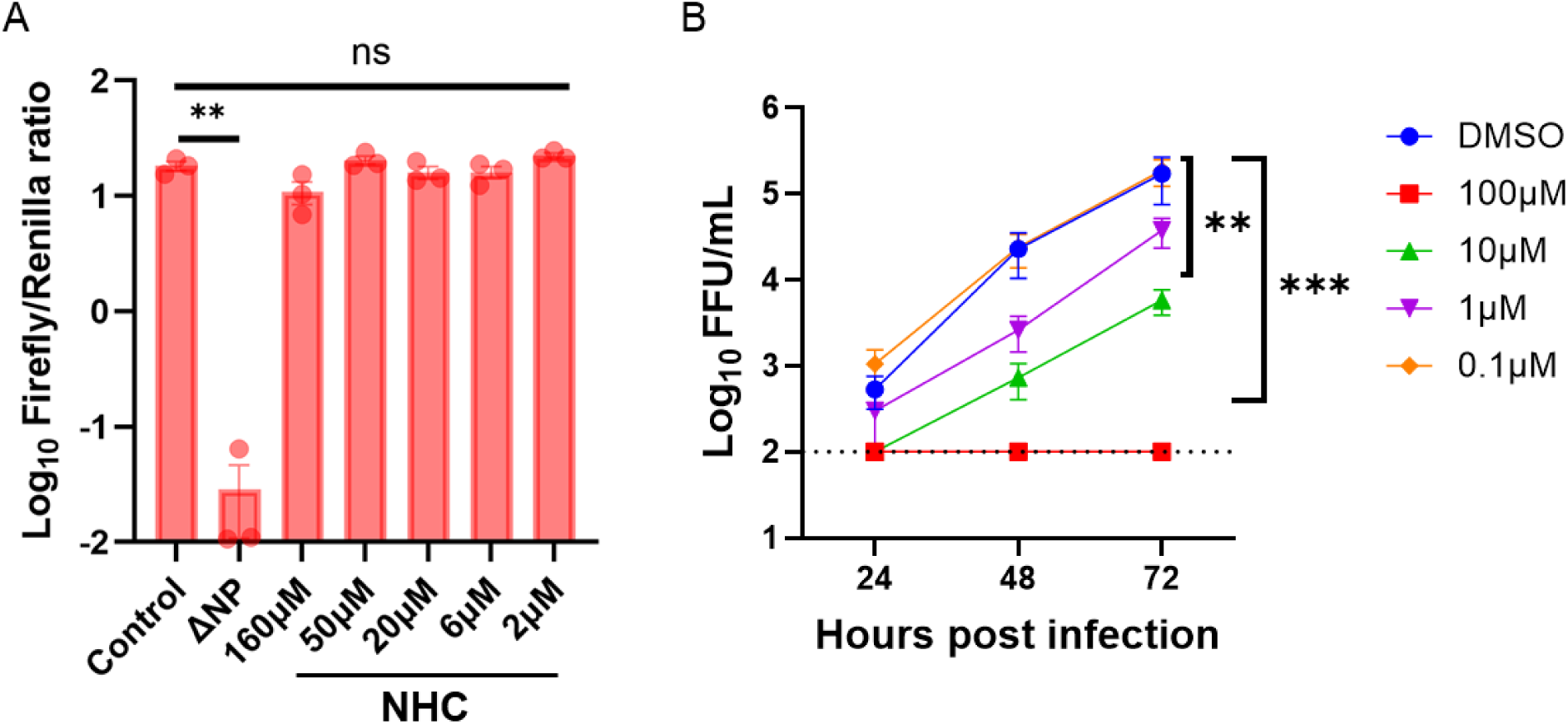
The antiviral mechanism of molnupiravir is independent of BRBV RdRp inhibition. (**A**) The effect of different concentrations of molnupiravir on the BRBV polymerase activity was quantified by using BRBV mini-genome reporter assay. Expression plasmids encoding for the BRBV polymerases (PB2, PB1, PA) and NP of BRBV and Renilla luciferase, plus the reporter construct (Firefly luciferase flanked by the 3’ and 5’ UTR of segment 2 of BRBV) were transfected into 293T cells in the presence or absence of different concentrations of molnupiravir (NHC). At 48 h post-transfection, the ratio of firefly luciferase to renilla luciferase activities was measured. Values are geomeans (± standard error of the geomean) of the luciferase activity from three experiments performed in duplicate. **, *P <* 0.01, by one-way ANOVA and Dunnett’s multiple comparisons. (**B**) 293T cells inoculated with BRBV (MOI of 0.01) in the presence or absence of different concentrations of molnupiravir. The virus titer in the culture supernatant was quantified in 24, 48, and 72 h. Values are means (± standard error of the mean) of the virus titer from three experiments performed in duplicate Data were analyzed by one-way ANOVA and Dunnett’s multiple comparisons on the area under the curves fit by linear regression of log-transformed virus titer over time. ** *P <* 0.01; *** *P <* 0.005. The dotted line represents the limit of detection at 200 PFU/mL.

### Molnupiravir inhibits Thogoto virus (THOV) infection *in vitro* and Dhori virus (DHOV) infection *in vitro* and *in vivo*

THOV, DHOV, and BRBV belong to the genus *Thogotovirus*. To assess the effects of molnupiravir on related viruses, we assessed its antiviral activity against THOV and DHOV replication in A549 cells (**Fig. 7A-B**). Like its effects on BRBV, molnupiravir reduced THOV and DHOV virus yield in the supernatants of treated cells at 48 hpi (**Fig. 7A-B**). Subsequently, we tested the *in vivo* efficacy of molnupiravir against DHOV; unlike BRBV, DHOV causes lethal infection in wild-type C57BL/6J mice [14, 26]. Male and female wild-type C57BL/6J mice received a high dose (500 mg/kg) of molnupiravir per oral gavage, 4 h prior to a lethal DHOV challenge (10X MLD_50_ (∼1pfu)). Mice continued to receive molnupiravir twice daily for 8 days and weight loss, clinical score, and survival of the mice were monitored for 10 days.

**Figure 7.**
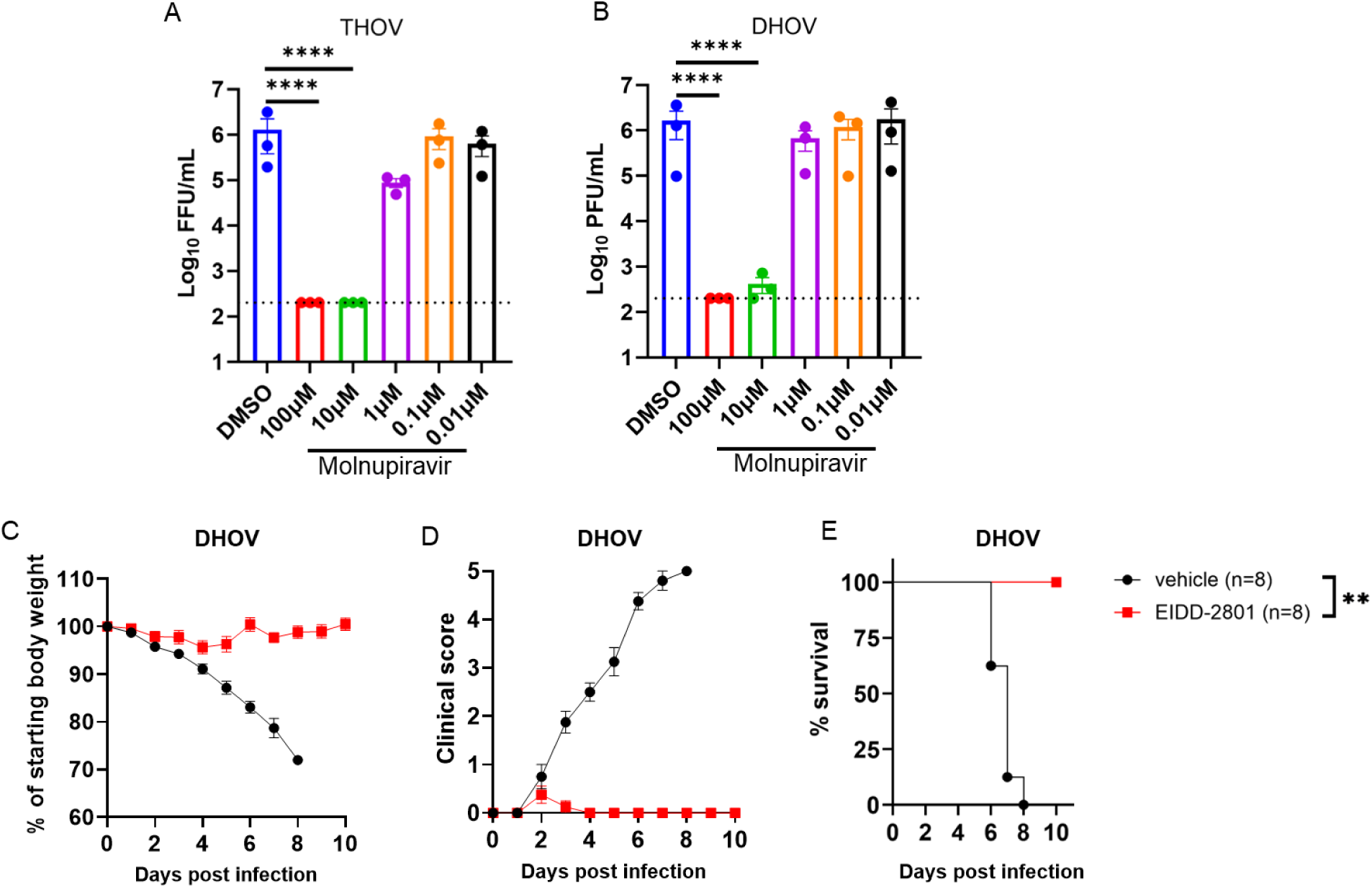
Molnupiravir suppresses THOV and DHOV infection in cell culture and protects wild-type C57BL/6 mice from lethal DHOV infection. (**A-B**) A549 cells inoculated with THOV and DHOV (MOI of 0.01) in the presence or absence of different concentrations of molnupiravir. Virus yield in the cell supernatants of THOV (**A**) and DHOV (**B**) at 48 hpi was measured and compared with control (DMSO). Values are means (± standard error of the mean) of the virus titer from three experiments performed in duplicate. ****, *P <* 0.0001, One-way ANOVA followed by Dunnett’s multiple comparisons. (**C-E**) C57BL/6J mice were treated twice daily with 500 mg/kg of molnupiravir for 8 days starting 4 h prior to intraperitoneal inoculation with 10X MLD_50_ of DHOV. Weight change (**C**), clinical score (**D**), and survival rates (**E**) were monitored for 10 days (n = 8/group, 2 independent experiments). Values are the means (± standard error of the mean) of the groups. Weight change and clinical scores are analyzed by the One-way ANOVA followed by Dunnett’s multiple comparison and survival by the log-rank test. **, *P <* 0.01.

Compared to vehicle control treated wild-type mice, molnupiravir-treated mice showed reduced weight loss (*P* < 0.05), clinical score (*P* < 0.01), and mortality (*P* < 0.01) after infection (**Fig. 7C-D**). Collectively, these data show that molnupiravir has antiviral activity *in vitro* and *in vivo* against multiple members of the *Thogotovirus* genus.

## DISCUSSION

Tick-transmitted BRBV [2, 10] and Oz virus (OZV) [27] are pathogenic viruses that can lead to severe and fatal infection and disease in humans. Moreover, seroepidemiological studies show that human infections with BRBV [11, 12], OZV [28], and other family members of the genus *Thogotovirus* (THOV, DHOV) [29] are more common than previously known. Currently, there is no approved therapy or vaccine against BRBV or related viruses. Here we report that BRBV replication is inhibited *in vitro* by the broad-spectrum antiviral drug molnupiravir. In mice, molnupiravir exhibited prophylactic and therapeutic activity against fatal BRBV disease. Moreover, we found that molnupiravir reduces viral titers and alleviates BRBV-associated pathology and clinical disease in mice. Finally, our data shows that the antiviral activity of molnupiravir extends to other family members of the Thogotovirus genus.

We tested several nucleoside analogs against BRBV that show antiviral activity against other RNA viruses. As BRBV is a member of the family of *Orthomyxoviridae*, like influenza A virus (IAV), we hypothesized that a compound that shows antiviral activity against IAV might show efficacy against BRBV. Previously, we showed that favipiravir, an antiviral with known activity against IAV, inhibits BRBV [10]; thus, we used it as a positive control. Of the four additional compounds included in our screening, two (galidesivir and molnupiravir) showed antiviral activity against IAV. Importantly, all three compounds were substantially more effective (lower EC_50_ values) against IAV compared to BRBV [30–32]. The basis for this difference is not known but may be due to structural differences between the polymerase complexes of the different viruses. The structure of the THOV RdRp complex was recently described and could be used to develop antiviral compounds with greater activity against BRBV and related viruses [33]. Nevertheless, it will be important to test future anti-influenza virus compounds against BRBV to identify additional possible inhibitors.

Molnupiravir was originally intended to treat alphavirus infections but was later tested for the treatment of IAV and showed efficacy in preclinical animal models [18, 34]. Subsequently, molnupiravir was tested as a treatment for SARS-CoV2 infection during the COVID-19 pandemic, leading to the 2023 FDA release of an EUA for molnupiravir as treatment of mild-to-moderate infection in adults at risk for progression to severe COVID-19. Additionally, preclinical testing of molnupiravir shows efficacy against Ebola, La Crosse, and Venezuelan equine encephalitis viruses [35–37]. In our current study, we showed that molnupiravir inhibits BRBV infection, emphasizing its application as a broad-spectrum antiviral agent that is licensed in the USA.

To better understand the antiviral mechanism of molnupiravir, researchers have performed various experiments. These studies showed that NHC (active metabolite of molnupiravir) plays a role in viral replication, promoting nucleotide G to A and C to U substitutions. This leads to mutations in viral synthesis, ultimately inactivating the resulting progeny viruses including influenza and SARS-CoV-2 [30, 38]. NHC can pair with either A or G in the viral genome which subsequently disrupts the typical base pairing G with C and A with U or T. In subsequent rounds of replication, mutations such as G to A (G: NHC: A) and C to U (C: G: NHC: A: U) are observed as NHC pairs with A or G. These mutations accumulate over time and exsert strong antiviral activity of molnupiravir [38]. In contrast, well-known RdRp inhibitors such as favipiravir and remdesivir, are incorporated into the viral genome and induce RNA chain termination or RNA stalling during replication respectively [39]. We observed that molnupiravir did not affect BRBV RdRp activity at 48 hours but inhibited virus production. Thus, these observations suggest that the anti-viral mechanism of molnupiravir against BRBV is due to the accumulated RNA mutations that induce error catastrophe.

Our study also extends the clinical and pathophysiologic characterization of our pre-clinical BRBV mouse model. Although many tick-borne viruses, including Severe Fever with Thrombocytopenia Syndrome (SFTS), Crimean Congo Hemorrhagic Fever (CCHF), Tick-Borne Encephalitis Virus (TBEV), and Heartland virus (HRTV), cause mild to severe thrombocytopenia [10, 40–42], the specific pathophysiologic mechanisms are often not well defined. Likewise, patients with BRBV infection develop acute thrombocytopenia, leukopenia, and lymphopenia [10], in conjunction with splenomegaly and liver injury that can progress to multiorgan failure and death in susceptible individuals. In this study, we observed that BRBV-infected *Ifnar1*^-/-^ mice developed severe leukopenia and thrombocytopenia, along with splenic enlargement and evidence of substantial spleen and liver injury, characteristics of severe disease in humans. Importantly, molnupiravir improved splenomegaly, splenic, and liver damage, as well as leukopenia and thrombocytopenia associated with BRBV infection, which likely contributed to the reduced disease in mice.

In conclusion, our study has shown that molnupiravir, an FDA-approved antiviral agent, can treat ongoing BRBV infection in a pre-clinical animal model. Further studies are warranted to explore the use of molnupiravir as a putative antiviral agent for the treatment of BRBV, and possibly related viruses, in humans.

## ACKNOWLEDGEMENTS

We thank Drs. Ishmael Aziati and Astha Joshi for providing technical assistance on BRBV RT-qPCR analysis.

## SUPPLEMENTARY DATA

**Supplementary Figure 1.**
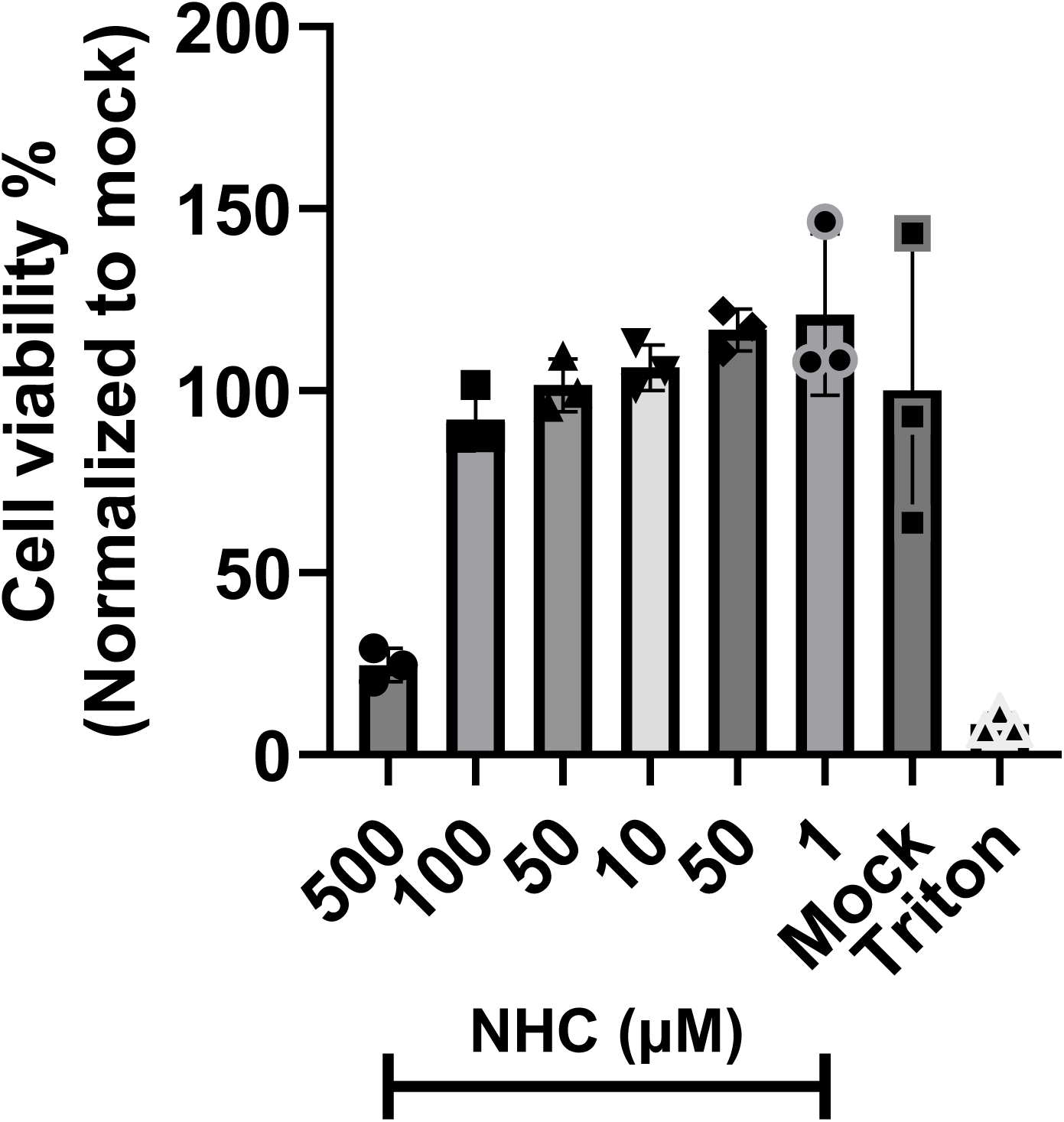
The viability of the A549 cells was minimally affected by molnupiravir. A549 cells were incubated with different concentrations of molnupiravir. Cell viability at 48 h was measured by using the Cell Titer Blue assay. DMSO was used as the mock control, and Triton X-100 was used as the positive control for cell death. Values are means (± standard error of the mean) of the cell viability from three experiments performed in duplicate.

**Supplementary Figure 2.**
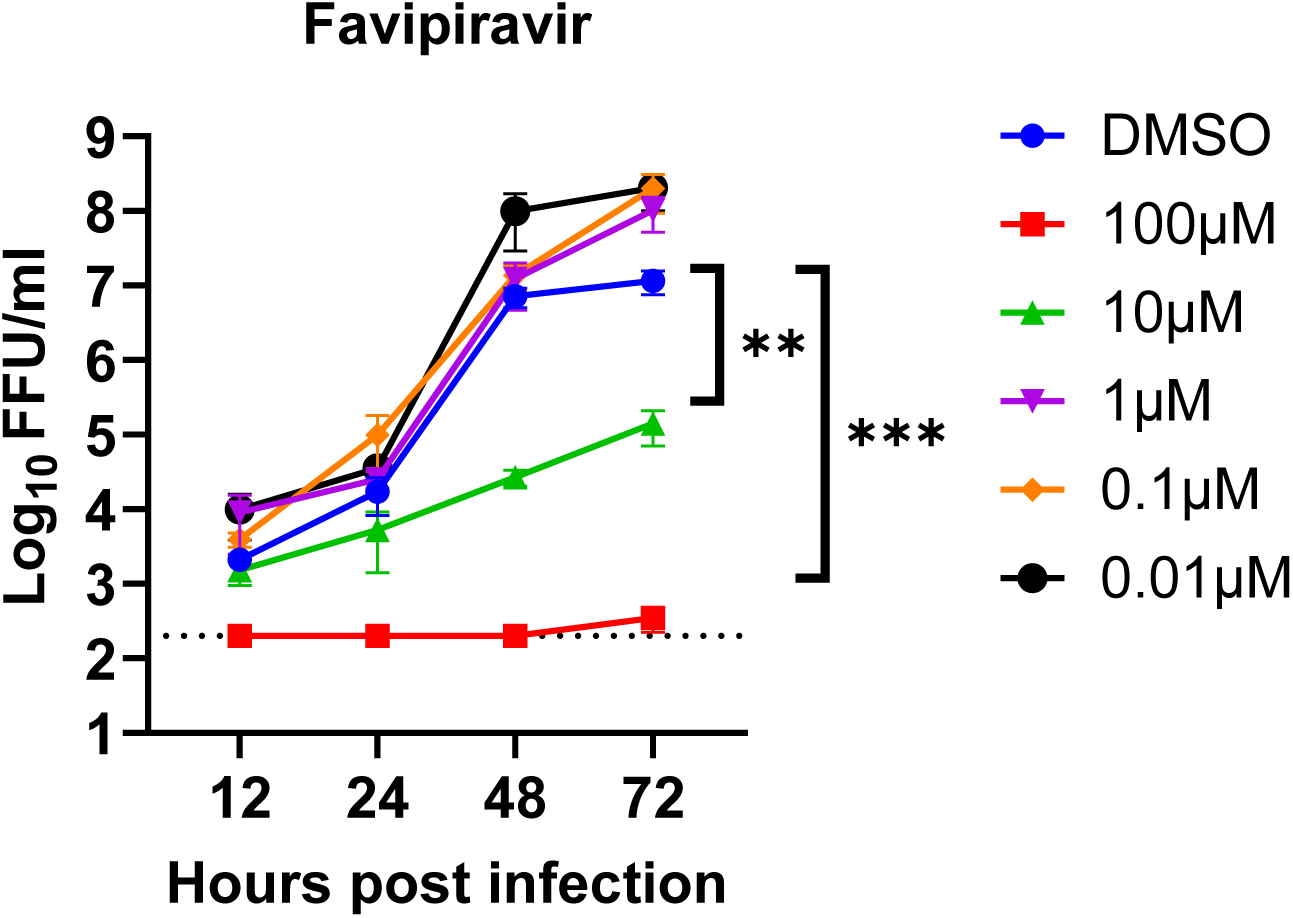
Favipiravir inhibits BRBV infection in cell culture. A549 cells inoculated with BRBV (MOI of 0.01) in the presence or absence of different concentrations of favipiravir. The virus titer in the culture supernatant was quantified at 12, 24, 48, and 72 h. Values are means (± standard error of the mean) of the virus titer from three experiments performed in duplicate. **, *P <* 0.01, *** *P <* 0.005 by one-way ANOVA on the area under the curves fit by linear regression of log-transformed virus titer over time and compared with Dunnett’s multiple comparisons. The dotted line represents the limit of detection at 200 FFU/mL.

**Supplementary Figure 3:**
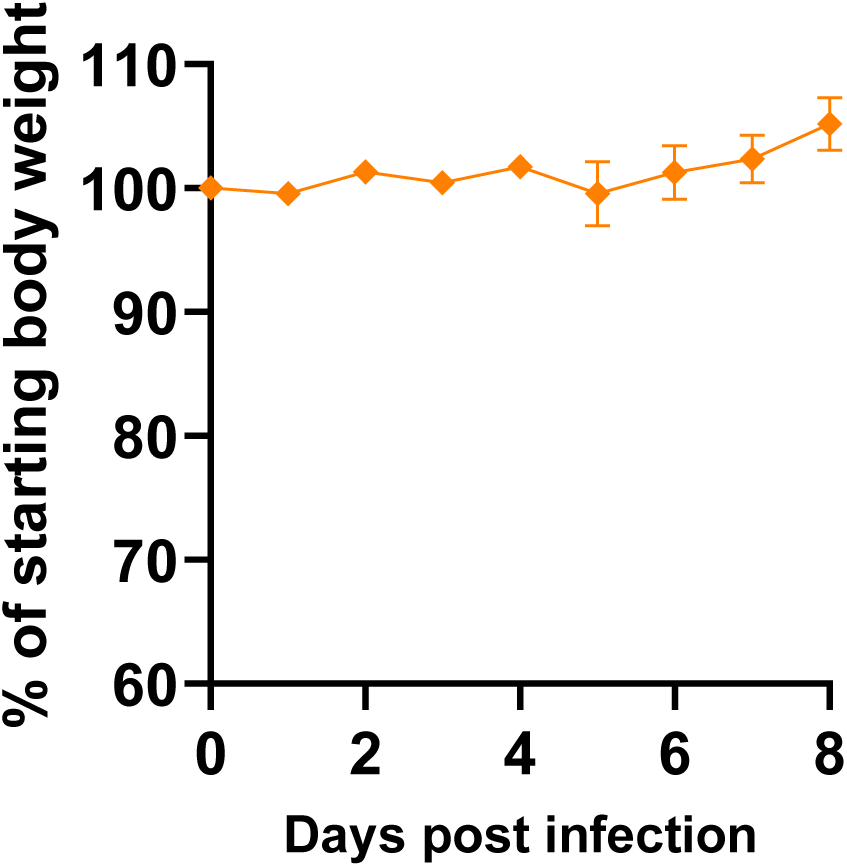
Molnupiravir (EIDD-2801) is well tolerated by *Ifnar1* ^-/-^ mice. Uninfected *Ifnar1*^-/-^ mice (n = 2) were treated orally with 150 mg/kg (mid dose) of molnupiravir twice daily for 8 days. Weight change was monitored for 8 days. Values are the means (± standard error of the mean) of the treated group.

**Supplementary Figure 4:**
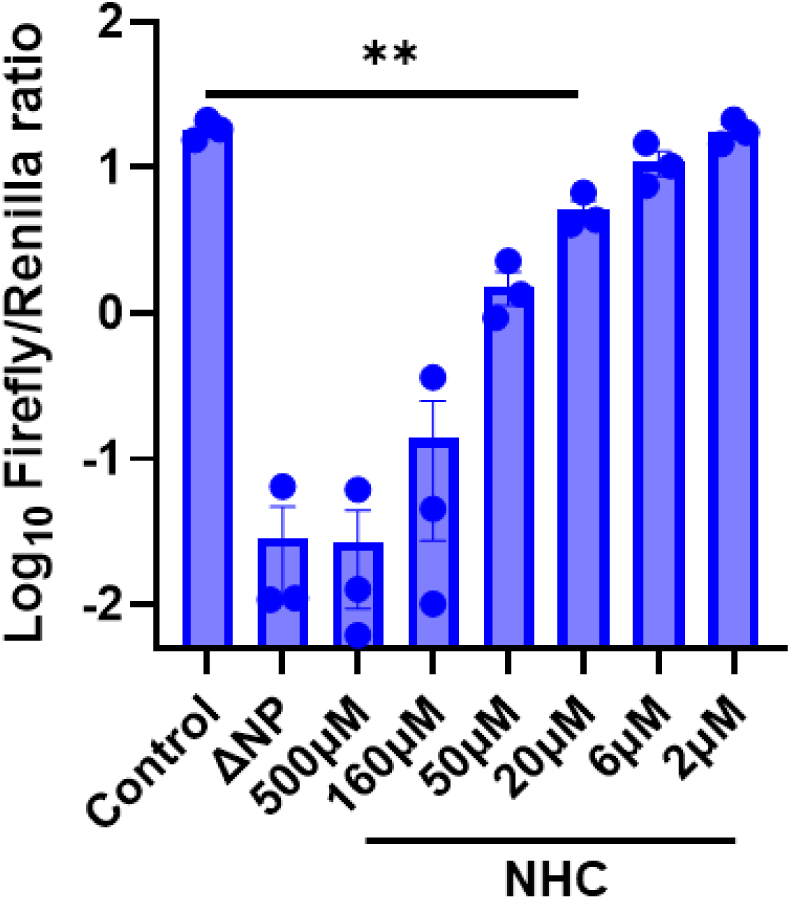
Favipiravir inhibits BRBV polymerase activity. The effect of different concentrations of favipiravir on the BRBV polymerase activity was quantified using the BRBV mini-genome assay. Expression plasmids encoding for the BRBV polymerases (PB2, PB1, PA) and NP of BRBV and Renilla luciferase, plus the reporter construct (Firefly luciferase flanked by the 3’ and 5’ UTR of segment 2 of BRBV) were transfected into 293T cells in the presence or absence of different concentrations of favipiravir. At 48 h post-transfection, the ratio of firefly luciferase to renilla luciferase activities was measured. Values are geomeans (± standard error of the geomean) of the luciferase activity from three experiments performed in duplicate. **, *P <* 0.01, by one-way ANOVA and Dunnett’s multiple comparisons.

**Supplementary Figure 5:**
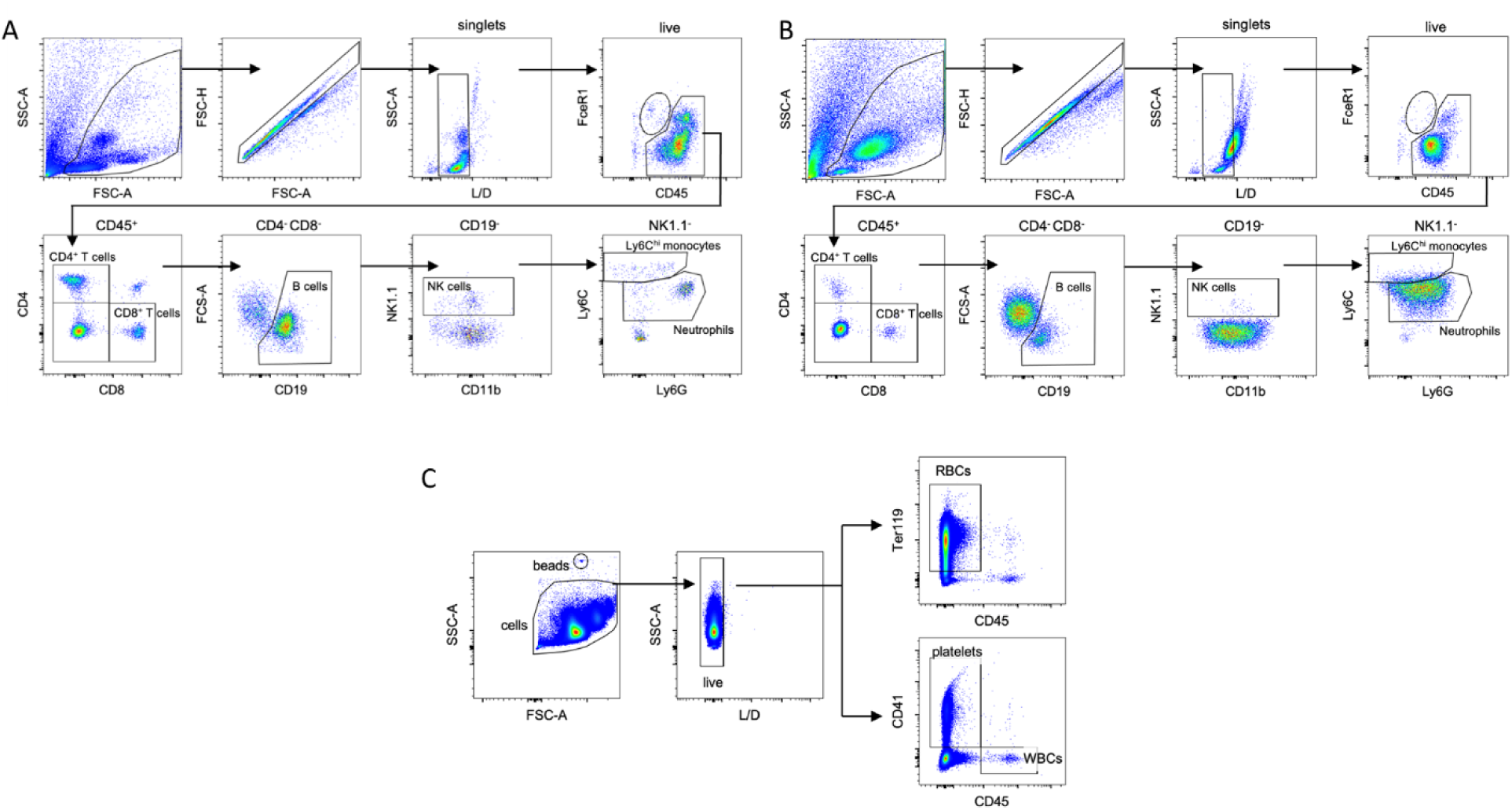
(**A-C**) Flow cytometry gating strategy for (**A**) uninfected spleen cells (**B**) infected spleen cells and (**C**) peripheral blood RBC, WBC, and platelets.

